# Uncovering expression signatures of synergistic drug response using an ensemble of explainable AI models

**DOI:** 10.1101/2021.10.06.463409

**Authors:** Joseph D. Janizek, Ayse B. Dincer, Safiye Celik, Hugh Chen, William Chen, Kamila Naxerova, Su-In Lee

**Affiliations:** Paul G. Allen School of Computer Science and Engineering, University of Washington; Medical Scientist Training Program, University of Washington; Recursion Pharmaceuticals; Center for Systems Biology, Massachusetts General Hospital and Harvard Medical School; Department of Radiology, Massachusetts General Hospital and Harvard Medical School

## Abstract

Complex machine learning models are poised to revolutionize the treatment of diseases like acute myeloid leukemia (AML) by helping physicians choose optimal combinations of anti-cancer drugs based on molecular features. While accurate predictions are important, it is equally important to be able to learn about the underlying molecular basis of anti-cancer drug synergy. Explainable AI (XAI) offers a promising new route for data-driven cancer pharmacology, combining highly accurate models with interpretable insights into model decisions. Due to the highly correlated, high-dimensional nature of cancer transcriptomic data, however, we find that existing XAI approaches are suboptimal when applied naively to large transcriptomic datasets. We show how a novel approach based on model ensembling helps to increase the quality of explanations. We then use our method to demonstrate that a hematopoietic differentiation signature underlies synergy for a variety of anti-AML drug combinations.

## 1 Main

Acute myeloid leukemia (AML) is the most commonly diagnosed form of leukemia in adults, and carries a poor prognosis [1]. While survival has improved over the past several decades for younger patients, older patients have not seen a similar improvement. This gap in survival has motivated the development of molecularly targeted combination therapies for patients who do not qualify for intensive induction chemotherapy [2]. Discovering optimal combinations of anti-cancer drugs is a difficult problem, however, as the space of all possible combinations of drugs and patients is large. While potentially synergistic drug combinations have traditionally been tested on the basis of either biological or clinical expert knowledge [3], more systematic approaches are necessary to effectively explore this space. Even systematic experimental approaches such as high-throughput screening are potentially insufficient, as there are hundreds of thousands of possible combinations of all anti-cancer drugs currently in development, each of which may have a different response in different patients [3, 4]. Therefore, predictive approaches are necessary to make the immense space of possible anti-cancer drug combinations manageable.

State-of-the-art predictive approaches fall short along another axis, however, by failing to provide biological insight into the molecular mechanisms underlying drug response, which is essential to facilitate the discovery of new and effective anti-cancer therapies [5–7]. While a wide variety of computational methods have historically been employed for drug combination prediction [8–12], recent work has demonstrated increased predictive performance using complex, non-linear machine learning (ML) models. For example, all of the winning teams in the AstraZeneca-Sanger Drug Combination Prediction DREAM Challenge utilized complex models in some part of their approach, including ensembles of random forest classifiers and gradient boosted machines [13]. Additionally, Preuer et al. have shown that deep neural networks outperform less sophisticated models such as linear models, achieving state-of-the-art performance at predicting the synergy of anti-cancer drug combinations in 39 cell lines [14]. A major weakness of these complex ML models is their “black box” nature; despite their high predictive accuracy, these models’ inner workings are opaque, making it challenging to gain mechanistic insights into the molecular basis of drug synergies. In cases where model interpretability is important, researchers resort to simpler, less accurate models like linear regression. For example, to identify genomic and transcriptomic markers associated with drug sensitivity, both the Cancer Genome Project [15] and the Cancer Cell Line Encyclopedia [16] used penalized elastic net regression.

Here, we present the EXPRESS (**ex**plainable **pr**edictions for gene expr**ess**ion data) framework to build both accurate *and* biologically interpretable ML models, addressing the accuracy-interpretability trade-off. A recent approach to manage this trade-off involves “explaining” complex predictive models using *feature attribution methods*, like Shapley values [17–20], to provide an importance score for each input feature (here, a gene). The Shapley value is a concept from game theory designed to fairly allocate credit to players in coalitional games [21]. By considering input features as players and the model’s output as the reward to be allocated [17], the most important features can be identified for complex models that would otherwise be uninterpretable. Unfortunately, the application of off-the-shelf feature attribution methods is unlikely to be successful in the context of large cancer ‘omics data. These methods are known to struggle in the setting of high-dimensional and highly correlated features, such as those present in transcriptome-wide gene expression measurements [22]. Furthermore, while complex ML models have been shown to achieve increased predictive performance when compared to simpler models, recent work has raised the concern that models with higher predictive performance do not necessarily have higher-quality attributions on the same tasks [23, 24]. Our results demonstrate how EXPRESS addresses these challenges.

First, using 240 synthetic datasets, we benchmark both classical and novel approaches and demonstrate how non-linearity and correlation in the data can impede the discovery of biologically relevant features. We then demonstrate that under conditions representative of typical biological applications, all existing approaches tend to perform poorly, and show how explaining ensembles of models improves the quality of feature attributions (Fig. 1a). Finally, we describe EXPRESS, which uses Shapley values to explain an *ensemble* of complex models trained to predict drug combination synergy on a dataset of 133 combinations of 46 anti-cancer drugs tested in *ex vivo* tumor samples from 285 patients with AML (Fig. 1b, Extended Data Fig. 1). In addition to building highly accurate predictive models, our ensemble interpretability approach identifies relevant biological signals underlying drug synergy patterns, most notably a gene expression signature related to hematopoietic differentiation. While individualized treatment for AML on the basis of cancer genomic signatures is already becoming an important aspect of clinical practice [6], our approach identifies a novel *expression*-based signature that is predictive of synergy across a broad class of drugs and their combinations in AML.

**Fig. 1.**
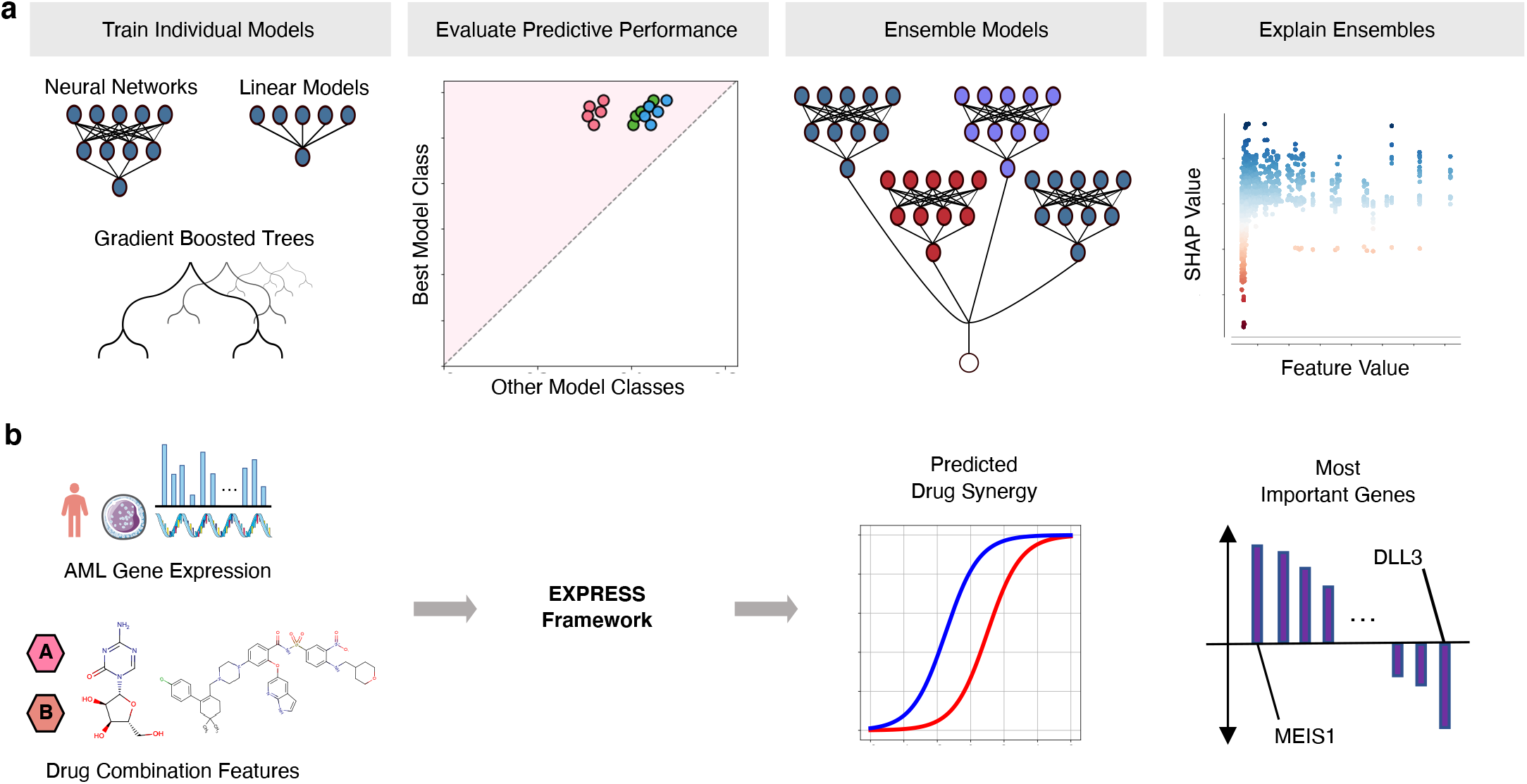
Overview of the study design. **a**, Our framework, EXPRESS (explainable predictions for gene expression data), for learning reliable explanations of cancer therapeutic machine learning models trained on high-dimensional gene expression data. After training a variety of individual models across multiple model classes, predictive performance is evaluated to select a best-performing model class. Multiple models from that class are then ensembled in order to produce more reliable and biologically-meaningful explanations. **b**, We apply our pipeline to a dataset of *ex vivo* anti-cancer drug synergy measurements for patients with acute myeloid leukemia, attaining not only superior prediction performance, but also identifying biological processes that are important for the determination of drug synergy.

## 2 Results

### 2.1 Current state-of-the-art explainable AI falls short on correlated features

Explainable AI (XAI) is a recent development in the ML community that attempts to provide a human-interpretable basis for the predictions of complex, “black box” models like neural networks. In particular, feature attribution methods are a class of methods that identify the relative importance of each input feature (e.g. the expression level of a gene) for a particular model [17, 25]. One popular feature attribution method involves applying Shapley values to interpret these complex models by measuring how much the model’s output changes on average when a feature is added to all other possible coalitions of features (Methods 4.1.1).

While applying explainable AI techniques to complex models has become a popular practice in the life sciences [26–31], applying these methods in the context of gene expression data is particularly difficult. Each patient will have a transcriptomic profile with tens of thousands of features with a high degree of feature interdependence (e.g., see the feature covariance matrix for AML transcriptomic data, Fig. 2, top right). This makes the task of accurate feature attribution harder for Shapley value algorithms, which ideally would operate on statistically independent features [17]. In the presence of correlated features, many models with diverse mechanisms could potentially fit the data equivalently well [32, 33]. Thus, even if we could explain a single model perfectly, that model might not correspond well to the true biological relationships between features and outcome.

**Fig. 2.**
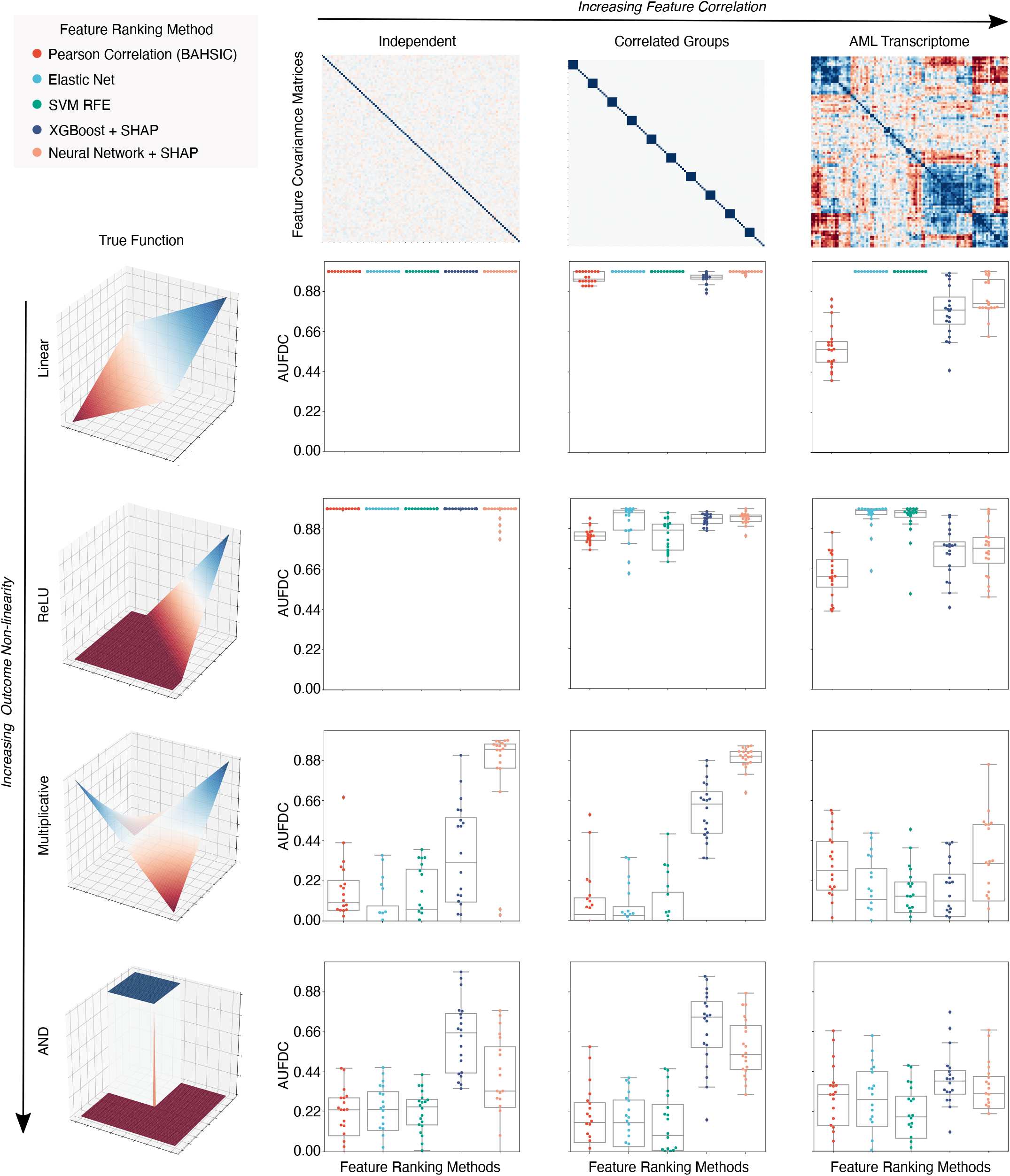
Benchmark metric reveals the impact of non-linearity and correlation on feature discovery. Each point in the box plots represents the benchmark score achieved by one of five feature ranking methods applied to one of 240 datasets generated from twelve synthetic or semi-synthetic dataset types (each subplot represents one dataset type). The rows (left) are sorted from top to bottom by increasing non-linearity of the true feature-outcome relationship (e.g. all datasets in the first row have a linear relationship between input features and outcome) while the columns are sorted from left to right by the increasing extent of the correlation between features in the dataset (e.g. all datasets in the last column have real AML bulk RNA-seq features). The metric plotted in each boxplot is the area under the feature discovery curve (AUFDC) (see Methods 4.2), where a higher score indicates better performance (0 represents random performance, while 1 represents perfect performance). For each dataset type (a pair of feature-outcome relationship and inter-feature correlation), 20 independent datasets are generated by randomly regenerating features. While all approaches achieve perfect performance on simple linear data with independent features (top left plot), all models have significantly worse performance as features become more correlated and outcomes become more non-linear (bottom right)

Since these conditions are ubiquitous in biological datasets, it is essential to understand how the efficacy of both Shapley value-based attributions and more conventional methods will be impacted in the setting of high-dimensional, highly correlated features. Measuring this efficacy is difficult, however, as existing benchmarks of feature attribution methods are designed to either measure the influence of features on the *particular model* being explained [18], or to measure the *predictive performance* of selected sets of features [34]. We therefore design a simple benchmark for this application (Fig. 2, Methods 4.2). To evaluate the effects of data correlation and non-linearity on feature attribution, we use 240 unique datasets. As input data, we consider synthetic datasets with independent features and synthetic datasets with multivariate normal covariance structure, as well as datasets with real gene expression measurements sampled from AML patients [7]. Since the goal of our benchmark is to define how well different methods recover *true features*, we create synthetic labels, allowing the ground truth to be recovered and measured. These labels are created by randomly sampling input features and relating them to the outcome using functions ranging from simple linear, univariate relationships to complex, non-linear step functions with interactions between features (Methods 4.2). For our metric of feature discovery performance, we measure how many *true features* are found cumulatively at each point in the lists of features ranked by each feature attribution method (Methods 4.2, Extended Data Fig. 2). Using this benchmark, we then evaluate five different methods for ranking biologically important features, including two complex machine learning methods (gradient boosted machines, neural networks) explained using Shapley values, as well as three more traditional linear methods: ranking features by their Pearson correlation with the outcome [35], ranking features by their elastic net coefficients [36], and recursive feature elimination using support vector machines [37].

When the outcome has a simple linear relationship with the input features, all approaches recover the true features well (see the perfect performance across all methods in the top left experiment in Fig. 2). When there is non-linearity in the data, however (see bottom two rows of Fig. 2), the complex machine learning models interpreted with Shapley values significantly outperform the linear approaches. Importantly, however, as the correlation between input features increases to the level seen in real AML transcriptomic data (Fig. 2, third column), all methods tend to perform poorly and there is a high degree of variance in the performance of each model class.

### 2.2 Ensembling overcomes variability in individual models

Given the observed variability of different models in terms of benchmark performance, a natural question that arises is how to select the predictive model that will attain the best performance at feature discovery. An intuitive solution is to simply pick the model with the best predictive performance. When we examine the relationship between predictive performance and feature discovery, however, we see that this is not necessarily a reliable strategy. For each of three popular model classes (linear models, feed-forward neural networks, and gradient boosted machines), we train twenty independent models on bootstrap resampled versions of the same dataset and measure test set prediction error and feature discovery performance. While there was significant overall correlation between test error and feature discovery, within each model class test error was *not* significantly correlated with feature discovery performance (see Fig. 3a,b). Therefore, while predictive performance may help to select a *model class*, it will not necessarily help to select which model within that class has the most biologically-relevant explanations.

**Fig. 3.**
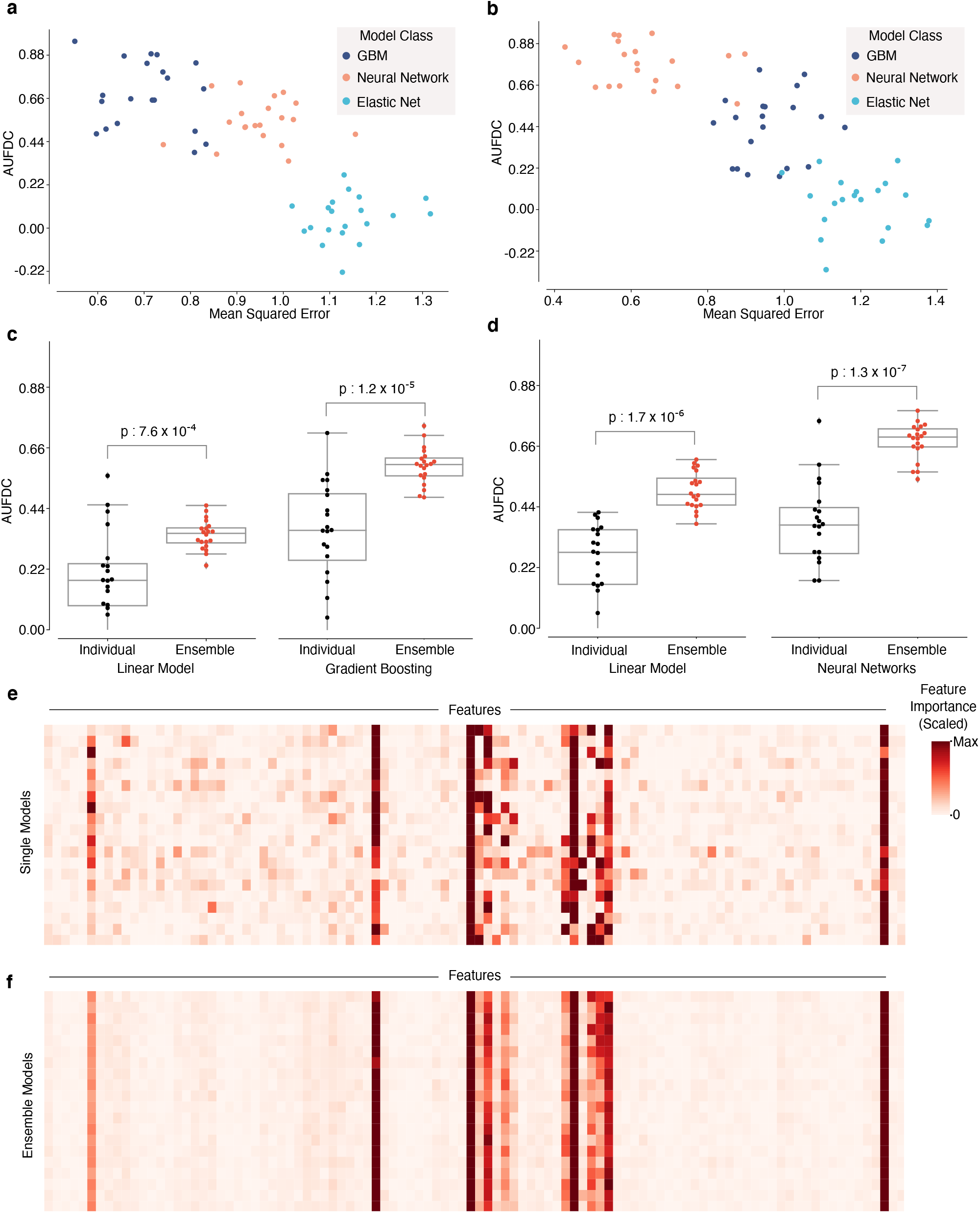
Explaining ensembles helps overcome instability in feature discovery performance for single models. **a-b**, Relationship between test error and feature discovery performance on bootstrap resampled versions of two synthetic datasets. Both datasets had clusters of highly-correlated features; one had a step function outcome (**a**) while the second had a multiplicative outcome (**b**). While there is a high overall correlation between test error and feature discovery performance for both datasets (Step Function Dataset: Pearson’s *r* = −0.77, *p* = 1.1 10^−12^; Multiplicative Dataset: Pearson’s *r* = −0.82, *p* = 1.2 10^−15^), there is no significant correlation after conditioning on model class (Elastic Net + Step Function Dataset: Pearson’s *r* = 0.19, *p* = 0.43; Neural Network + Step Function Dataset: Pearson’s *r* = 0.02, *p* = 0.94; XGBoost + Step Function Dataset: Pearson’s *r* = −0.18, *p* = 0.45; Elastic Net + Multiplicative Dataset: Pearson’s *r* = −0.11, *p* = 0.65; Neural Network + Multiplicative Dataset: Pearson’s *r* = −0.22, *p* = 0.35; XGBoost + Multiplicative Dataset: Pearson’s *r* = −0.13, *p* = 0.60;). **c-d**, Comparison of feature discovery performance between individual models and ensemble models on the AML gene expression dataset with a step function outcome (**c**) and a multiplicative outcome (**d**). **e**, Heatmap of feature attributions for 20 individual XGBoost models trained on bootstrap resampled versions of the correlated features Step Function dataset. **f**, Heatmap of feature attributions for 20 ensembles of XGBoost models trained on bootstrap resampled versions of the correlated features Step Function dataset.

Furthermore, when we examine the feature attributions across individual models within a single model class, we observe that they vary significantly from model to model (Fig. 3e). This indicates a lack of stability in the attributions: minor perturbations to the training set (such as bootstrap resampling) can lead to significant variability in the features identified as most important by the model [32], and prior work in machine learning applied to human genomics and epigenomics has suggested the necessity of considering multiple models when analyzing explanations [38, 39]. Likewise, recent work on feature selection for black box predictive models in healthcare has pointed out the need to select robust features [40].

While ensembling machine learning models is classically known to increase the accuracy of models by increasing stability of predictors, it remains to be demonstrated whether ensembling can improve biological hypothesis generation. We therefore created ensembles of models for the datasets with the worst performance in the original benchmark task – the datasets with patient AML features and non-linear outcomes, and found that ensembling not only decreases the variance in feature discovery performance, but also significantly increases the average feature discovery performance of the ensemble models (Fig. 3c,d). We observe a similar result in the datasets with highly correlated Gaussian features (Extended Data Fig. 3).

To understand how the ensembled models differed from the individual models, we analyzed the difference between the attributions attained by a variety of ensembled models and the individual models. We see that the variability in attributions across bootstrap resampled versions of the dataset significantly decreases (Fig. 3f, Extended Data Fig. 4). Furthermore, when compared to the single models, the ensemble models tend to place more weight on a small set of important features and attribute less importance to spurious correlates: spurious correlations cancel out over repeated model trainings, while true signal remains consistent (Extended Data Fig. 4).

**Fig. 4.**
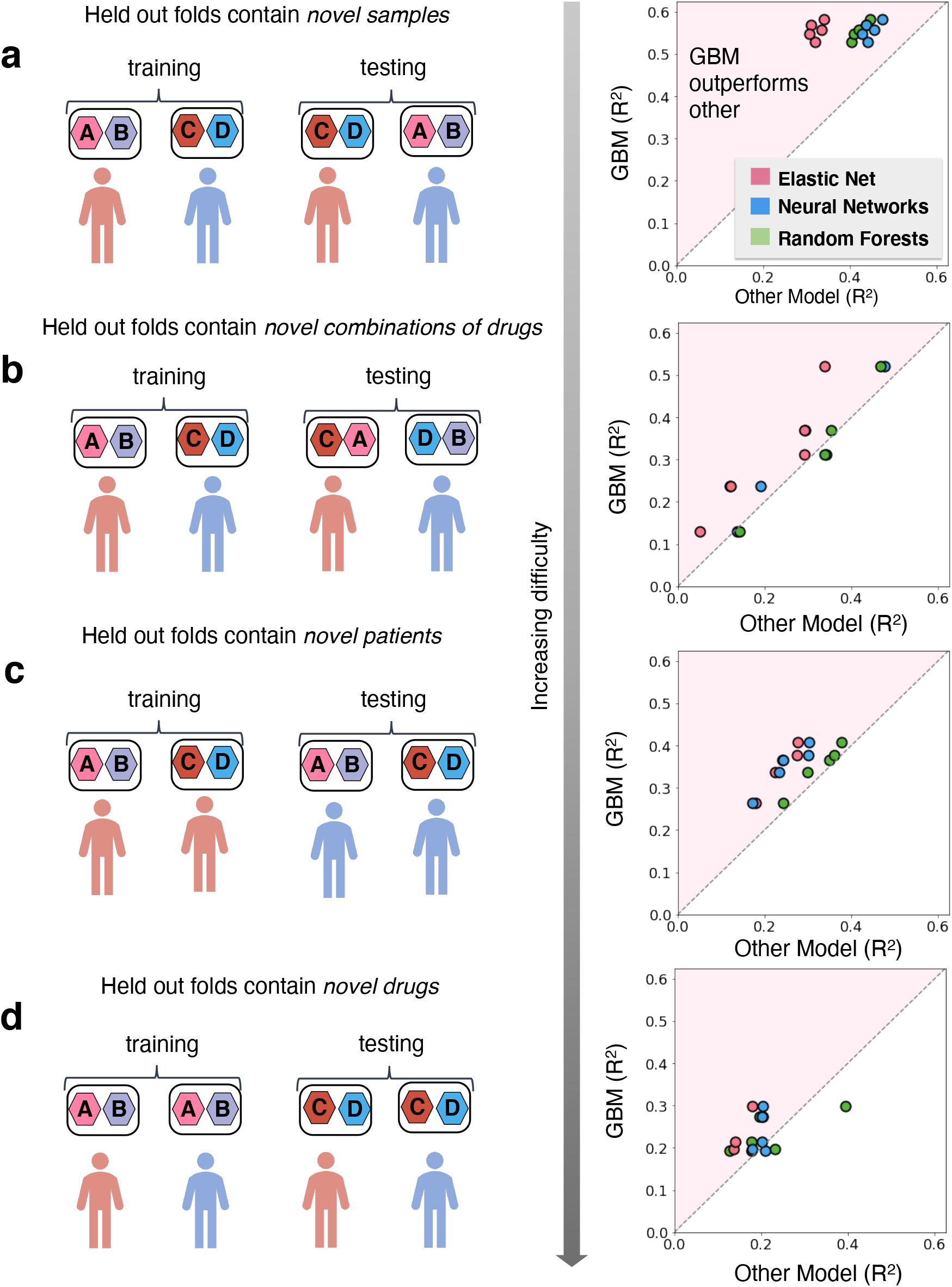
Comparison of predictive performance between model classes across four stratification settings. Each point in the plots on the right represents an evaluation of model performance after a different split of the data. In order to consider a variety of potentially useful application settings, samples were stratified in four ways. Each sample comprises primary tumor cells from a patient with AML and a pair of anti-cancer drugs. In **a**, samples are randomly split into 5 different train test folds. In **b**, samples are split on the basis of the drug combinations, so that held out test folds contain novel drug combinations not present in the training data. In **c**, samples are split on the basis of patients, so that held out test folds contain patients not present in the training data. In **d**, samples are split on the basis of individual drugs, so that held out folds contain drugs not present in the training data.

These results suggest a natural approach for applying explainable AI techniques to complex biological datasets (see Fig. 1a). A variety of model classes should be compared in terms of predictive performance, and following the selection of the best performing model class, the set of well-performing models from that class should be ensembled for explanation.

### 2.3 Complex gradient boosted machines accurately predict drug synergy in AML samples

After determining the importance of model class selection and model ensembling from our benchmark, we applied our framework to publicly available data provided by the Beat AML collaboration [7]. These data consist of the gene expression profiles of primary tumor cells from 285 patients with acute myeloid leukemia, as well as drug synergy measured for these cells in an *ex vivo* sensitivity assay for 131 pairs of 46 distinct drugs, spanning a variety of cancer subtypes and anti-cancer drug classes (Extended Data Fig. 1 and Supplemental Tables 1 and 2). The input features of each sample thus comprise ‘gene expression features’ which describe the corresponding patient’s tumor’s molecular profile, and ‘drug features’ which describe the two drugs in that combination in terms of the gene targets of each of the two drugs (Fig. 1b and Methods).

EXPRESS begins by comparing multiple model classes: elastic net [36], deep neural networks [41], random forests [42], and extreme gradient boosting (XGBoost) [43], in terms of the test error calculated using 5-fold cross validation tests. To rigorously evaluate the predictive performance of the models, we performed comparisons using four different schemes for stratifying samples into train and test sets. Each different stratification assesses the generalization performance for a different possible application scenario (see Fig. 4 and Methods) [14, 44]. Across these four settings, XGBoost shows better performance in 53 comparisons out of 60 (=4×3×5) comparisons from four settings, with three alternative methods, and for five test folds. Elastic net, random forests, and deep neural networks show better performance in 4, 27, and 30 comparisons, respectively. Our framework therefore selects XGBoost as the optimal model class for further downstream interpretive analysis.

### 2.4 Ensembled attributions reveal important genes for anti-AML drug synergy

After identifying gradient boosted machines as the best-performing model class for our dataset, we ensembled individual models until the ensemble model attributions were stable, leading to a final ensemble of 100 XGBoost models (see Extended Data Fig. 5, Methods). We then analyzed the resultant ensemble model attributions to look for genes with *global* importance for drug combination synergy, i.e., genes whose expression is related to synergy across many different drug pairs in our dataset [18]. Genes that impact global synergy could belong to pathways with outsize importance to cancer biology which are targeted by many drugs in the dataset, such as MAPK signaling or PI3K-Akt signaling (see Supplemental Table 2 for a list of targets for each drug), or could be related to larger-scale transcriptional changes impacting many pathways simultaneously, such as the degree of differentiation of leukemic cells [45].

We first visualize genes with monotonic relationships with synergy across all samples in the dataset by plotting these robust attributions in a dependence plot. For example, a strong positive correlation between the expression level of MEIS1 (the 2nd strongest relationship out of 15,377 genes tested), and its attribution value indicates that patients with higher levels of MEIS1 expression are predicted to respond more synergistically to the drug pairs tested in this dataset (Fig. 5a). MEIS1 has been shown to be upregulated in mixed-lineage leukemia (MLL)-rearranged AML [46], while also driving leukemogenesis independently of MLL-rearrangement [47]. Recently, high MEIS1 expression has been observed within Venetoclax-resistant AML subclones with “monocytic” characteristics [48]. Because AML in different patients may manifest in different developmental stages [48], the importance of MEIS1 suggests that our model may be learning a differentiation-related expression signature underlying the synergistic ability of certain drugs to overcome resistance to others.

**Fig. 5.**
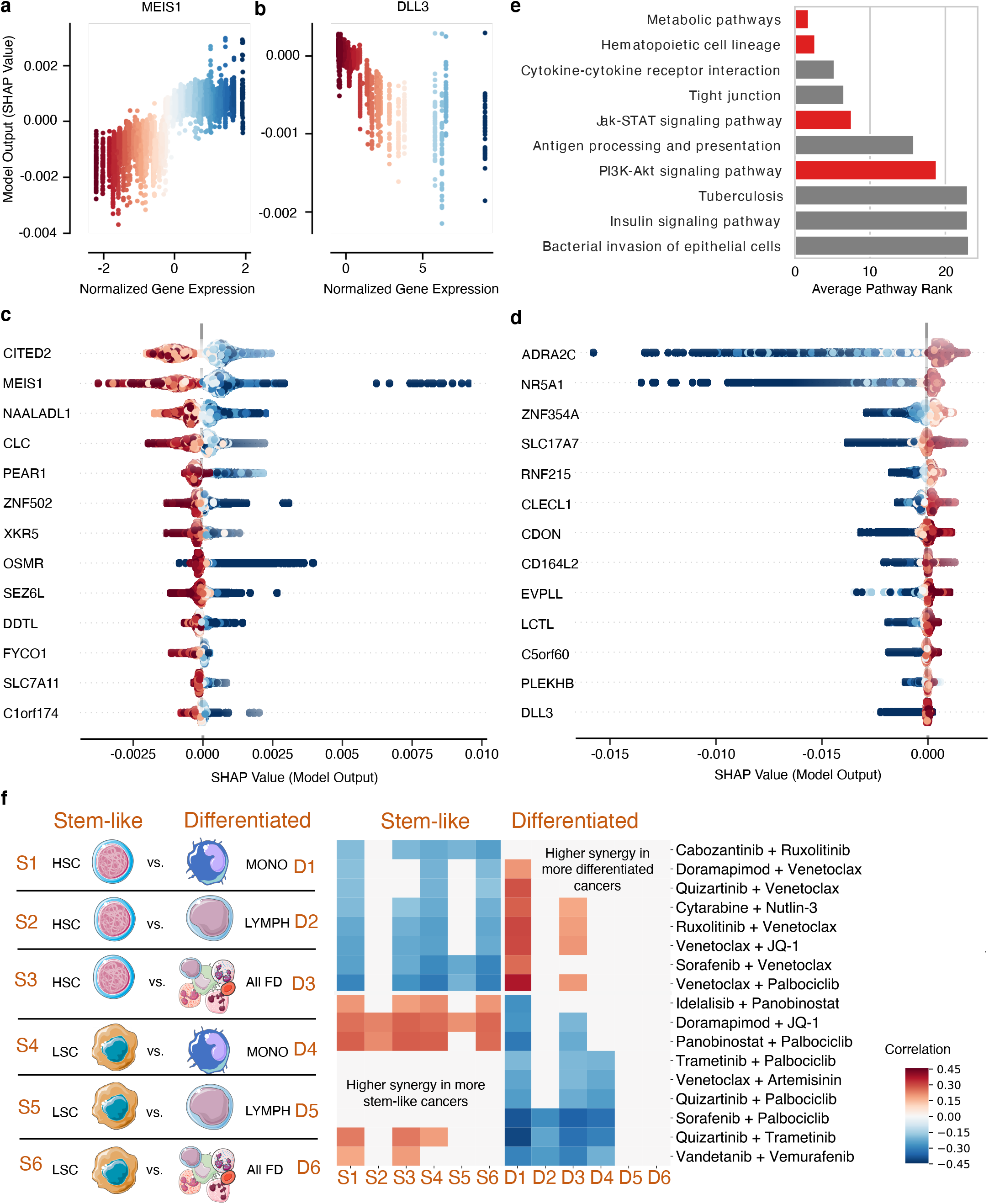
Transcriptomic factors affecting anti-AML drug combination synergy. **a-b**, SHAP dependency plots for MEIS1 and DLL3. Each point represents a single sample (one patient with a pair of anti-cancer drugs), the x-axis and color encoding represent the normalized gene expression values, while the y-axis represents the feature attribution value (change in predicted drug synergy attributable to that feature). **c-d**, SHAP summary plots for the transcripts with the strongest positive (**c**) and negative (**d**) relationships with anti-AML drug synergy.Each point still represents a single sample and the color encoding still represents normalized gene expression values, while the x-axis now represents the feature attribution value (plotted on the y-axis in the corresponding analysis in **a** and **b**). **e**, Biological pathways most highly enriched in the list of most important gene expression features, sorted by their average ranking across several top gene thresholds. **f**, For twelve separate differential gene expression profiles created by pairing gene expression measurements from a more stem-like hematopoietic lineage cell population (HSCs or LSCs) with a more differentiated hematopoietic lineage cell population (monocytes, lymphocytes, or all fully differentiated cells), we measured the correlation between the average expression of that profile and the synergy for each drug combination. After FDR correction, we plotted all combinations with significant correlations across at least two profiles. We find that some combinations of drugs tend to have higher synergy in more differentiated cancers, while some combinations of drugs tend to have higher synergy in more stem-like cancers.

EXPRESS can identify other genes showing such trends and visualize many of these feature attribution relationships at once by assembling the marginal distributions of the expression-attribution dependence plots into a summary plot. Figure 5c-d shows two summary plots – one for the genes where higher expression correlates with higher predicted synergy, and another for the genes with negatively correlated relationships (see Supplemental Table 3 for an exhaustive list). One of the top negatively correlated genes was DLL3 (Fig. 5b), a member of the Notch signaling pathway, which has been shown to have prognostic significance in patients with AML: patients with higher DLL3 expression have been shown to have lower overall survival [49]. We find that many of the top genes underlying synergy in both directions have been related to different stages of hematopoietic development. For example, CITED2 (the top positively correlated gene) is known to be essential for the maintenance of adult hematopoietic stem cells [50]. Additionally, CITED2-mediated hematopoietic stem cell maintenance has also been shown to be critical for the maintenance of AML [51]. Other genes in this list, such as OSMR, have further been shown to be essential for the maintenance of normal hematopoiesis [52]. Still other top genes, like SLC7A11 and SLC17A7, have been linked to prognosis of AML [53–55].

In addition to considering genes whose expression consistently impacts synergy either positively or negatively across all drug combinations, we additionally ranked genes by the magnitude of their global attribution values. This analysis allows genes that are important for multiple combinations to be ranked highly, even if higher expression of these genes are linked with higher synergy for some combinations and lower synergy for other combinations. When EXPRESS ranks all genes by the magnitude of their global attribution values (see Methods, Supplemental Table 4), we again find genes that are related to hematopoietic development and AML prognosis. In particular, IL-4 (top ranked gene by non-directional magnitude) is an important cytokine regulating the tumor microenvironment that has been shown to be specifically downregulated in AML compared to normal myeloid cells [56]. STAT6 (ranked 31st) is a transcriptional regulator known to be a key mediator of cytokine signaling [57]. It has previously been experimentally demonstrated using CRISPR-Cas9 genomic engineering that STAT6 specifically mediates IL-4-induced apoptosis in AML [58]. Furthermore, expression of STAT6 has been shown to be high in hematopoietic stem cells, but not in more differentiated progenitors [59]. Other top genes in this list, such as SLC51A and RNF213 (ranked 2nd and 6th overall, respectively), have been previously linked to AML and familial myelodysplasia via GWAS studies [60, 61].

### 2.5 Pathway explanations identify global importance of a differentiation signature

While attributions and trends for individual genes are informative, to gain systems-level insights into the processes important to drug synergy prediction, we can also use pathway databases to systematically check if genes from certain pathways are over-represented in EXPRESS’s top-ranked genes. When we test the top-ranked genes for pathway enrichment, we find that the top pathway (Fig. 5e) is related to cellular metabolism. Expression programs regulating cancer metabolism have previously been linked to resistance to a variety of the drugs tested in this dataset. For example, AML cells that are resistant to the tyrosine kinase inhibitor Cabozantinib have been shown to have higher glucose uptake, GAPDH activity, and lactate production than Cabozantinib-sensitive cells [62].

Furthermore, consistent with our hypothesis that the importance of MEIS1 for synergy may be linked to a differentiation signature, the second most highly enriched pathway contains genes that control differentiation along the hematopoietic cell lineage (*p* = 8.2 ×10^−3^). Previous studies have shown that leukemic stem cell signatures associate with worse clinical outcomes [63, 64], and cells at different differentiation stages have been shown to respond differently to particular combination therapies [65, 66]. The differentiation signature and metabolic signature may in fact be related, as prior work has shown that less differentiated leukemic cells have unique metabolic dependencies [67], and have even proposed metabolic changes as a mechanism mediating anti-cancer drug combination resistance specifically in stem-like leukemic cells [68].

To further validate the importance of differentiation signatures as a global pattern underlying drug combination synergy, we used an independent set of RNA-sequencing data generated from specific sub-populations of hematopoietic cells to create gene lists that are relatively more (or less) expressed in either hematopoietic stem cells (HSCs) or leukemic stem cells (LSCs) compared to more differentiated populations, such as monocytes, lymphocytes, and all fully differentiated blood cells (Methods) [69]. Considering six pairs of cell types (Fig. 5f, left) leads to 12 gene lists; six of the genes lists represent more stem-like expression states, while the other six lists represent more differentiated signatures (Methods section for more details). For each gene list, we measured the correlation between the average expression of the genes in the list and drug synergy for each drug pair (Fig. 5f, right), and plotted correlations that were significant after multiple hypothesis testing correction.

Remarkably, we found two distinct sets of drug combinations – combinations that were more synergistic when applied to tumor samples with more stem-like expression profiles, and combinations that were more synergistic when applied to tumor samples with more differentiated expression profiles (Fig. 5f, right). For instance, many combinations containing the BCL-2 inhibitor Venetoclax were associated with increased synergy when a more differentiated signature was present. Specifically, these were most strongly associated with a monocytic expression signature (Signature D1). Recent studies have demonstrated that in some patients, AML subclones with a monocytic differentiation signature exist next to subclones with a more primitive, stem-like transcriptional profile [48, 66]. Monocytic subclones have been shown to be relatively resistant to Venetoclax [48, 66], raising the possibility that the drugs paired with Venetoclax in the identified combinations could be helping to overcome this resistance. For example, our approach identifies the combination of Ruxolitinib, a JAK inhibitor, with Venetoclax as having more synergy in more differentiated cancers. The capacity of Ruxolitinib to synergize with Venetoclax, specifically by targeting and overcoming monocytic resistance, has recently been demonstrated in several studies [66, 70]. EXPRESS identifies a number of additional drugs that may be combined with Venetoclax to the same effect, including the p38 MAP kinase inhibitor Doramapimod, the tyrosine kinase inhibitors Quizartinib and Sorafenib, the cyclin-dependent kinase 4/6 inhibitor Palbociclib, and the BET bromodomain inhibitor JQ-1. Interestingly, EXPRESS also identifies a handful of combinations not containing Venetoclax for which synergy is also associated with a differentiation signature, including the combination of Cabozantinib and Ruxolitinib, as well as the combination of MDM2 inhibitor Nutlin-3 and the chemotherapeutic cytosine analogue Cytarabine.

These results show that the exact position of AML cells on a hematopoietic differentiation spectrum predicts the synergy that can be achieved with specific therapy combinations. Assessment of an AML stemness (or differentiation) signature may therefore be useful in guiding therapy choices in the clinic.

### 2.6 Feature interactions identify drug-specific gene expression signatures

In addition to identifying expression signatures that are generally relevant for drug synergy across many combinations, our approach is also able to identify genes and pathways that are relevant for *specific* drugs. To quantify these drug-specific mechanisms, we used an extension of the Shapley value called the Shapley interaction index [18, 71], which extends attributions for single features to interactions between pairs of features (see Methods). Intuitively, expression of a particular gene may be more important when one of the drugs in a combination is specifically targeting that gene. Likewise, expression of a particular gene may be less important when neither drug targets that gene. Therefore, to quantify which genes were important for specific drugs, we measured the interaction values between each drug feature label and all gene features.

By analyzing the most important genes for each drug ranked by the average magnitude of their interactions (see Methods), EXPRESS is capable of revealing the specific biological processes related to synergy for a particular drug. After generating interaction values between all genes and drugs, we tested each list of global drug-specific gene attributions for pathway enrichment (see Methods). We found that these enrichments aligned with prior knowledge of the mechanisms of the drugs in question (Fig. 6).

**Fig. 6.**
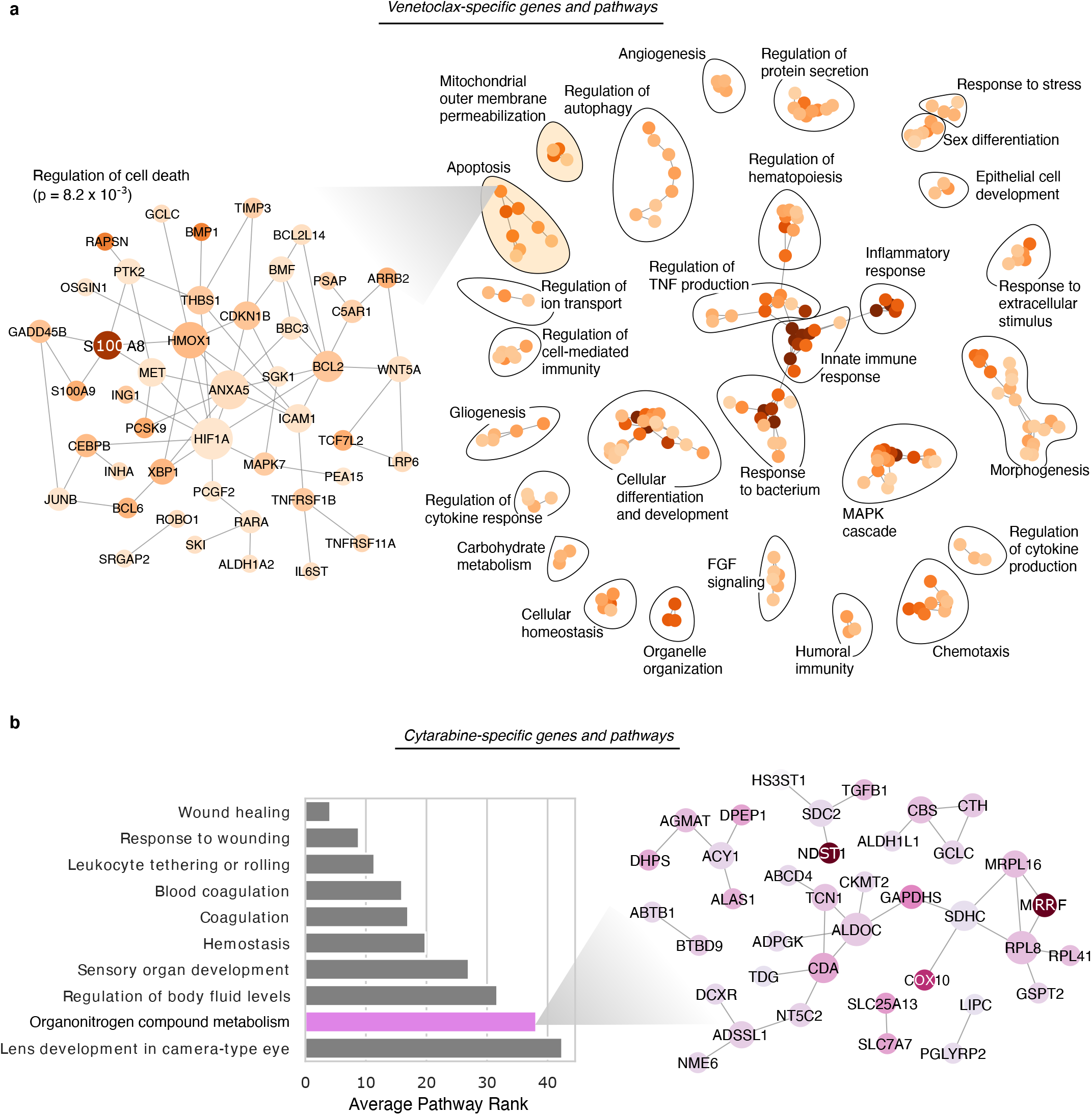
Transcriptomic factors affecting synergy of combinations including specific drugs. **a**, The top pathway enrichments in the set of transcripts affecting synergy of drug combinations including the drug Venetoclax. Each node in the graph on the right represents a single pathway, where the color indicates the strength of the enrichment, while edges indicate significant overlap in terms of the set of genes in each pathway. The zoomed inset graph on the left shows the genes in one pathway, “Regulation of cell death,” from the cluster of apoptosis-related pathways. In the left inset graph, each node is a gene, while the edges represent known protein-protein interactions. **b**, The top pathway enrichments in the set of transcripts affecting synergy of drug combinations including the drug Cytarabine. The bar plot (left) shows the top pathways, while the inset graph on the right shows the relevant genes from one pathway, “Organonitrogen compound metabolism.”

For instance, EXPRESS pinpoints genes involved in apoptosis as important determinants of synergy for pairs of drugs containing Venetoclax (Fig. 6a, right), a drug which functions by restoring apoptotic function in malignant cells via inhibition of the gene B Cell Lymphoma-2 (BCL-2) [72]. Examining the individual genes in one of the enriched pathway modules for Venetoclax (Fig. 6a left, “Regulation of cell death” term, FDR-corrected *p* = 8.0 × 10^−3^) reveals Venetoclax’s specific target BCL-2 to be an important predictive gene. Measuring the strength and direction of the relationship between BCL-2 expression and Venetoclax-specific BCL2 attribution values, we find that increased BCL2 expression is associated with markedly increased drug synergy in the context of Venetoclax treatment (Spearman *ρ* = 0.156, *p* = 4.0 ×10^−70^). Other genes in this module include S100A8 and S100A9, both genes which have previously been linked to patient response to Venetoclax as well as differential expression in hematopoietic stem cells compared to more differentiated populations [73, 74]. Other important biological processes detected by the Venetoclax-specific attributions include the MAPK cascade, which has been linked to Venetoclax-resistance through stabilization of MCL1 [75], and Fibroblast growth factor (FGF) signaling (see Fig 6a, right). Interestingly, Fibroblast Growth Factor 2 (FGF2) release by dying cells has recently been implicated as a transient, non-heritable mechanism of Venetoclax resistance [76], highlighting the power of transcriptomic analysis to discover phenomena not observable in mutational data alone.

As another example, one of the most enriched biological process terms for cytarabine, an organonitrogen compound, is the “metabolism of organonitrogen compounds” (Fig. 6b, *p* = 4.0 × 10^−5^). The individual genes in this module include CDA and NT5C2, two genes responsible for the metabolism of cytarabine that have previously been shown to be important genetic factors determining the response to cytarabine therapy [77]. We conducted the interaction drug-specific feature attribution analysis and pathway enrichment characterization for all drugs, which can serve as a resource for researchers interested in the particular mechanisms underlying AML response to these drugs (Supplemental Table 5). This analysis demonstrates that EXPRESS is able to identify not only expression trends important for large sets of combinations, but also for specific drugs.

## 3 Discussion

By ensembling complex models, the EXPRESS framework not only enables accurate predictive performance, but also robust and biologically meaningful explanations. While prior work has been able to attain high accuracy with complex models [14], our approach can provide explanations to assure patients, clinicians and scientists of the biological soundness of our predictions, even when models have high-dimensional input features with a high degree of feature correlation. The importance of interpretability in the context of biomedical AI is increasingly being recognized. Model explanations can help identify when apparently accurate “black box” models may in fact be relying upon unreliable confounders, also known as “shortcuts” [78, 79]. Explanations also allow physicians to communicate the logic of algorithmic decisions with patients, which can increase patient trust in the treatment process [80]. Finally, by displaying the logic underlying model decisions, explainable AI can enable better collaboration between physicians and AI models. For example, when applied to the Beat AML dataset [7], our model was optimized without respect to the cost or FDA approval status of different drug combinations. Where a “black box” model can only provide physicians with a synergy score for drug combinations, the mechanistic explanations provided by our model could help a physician to choose combinations with a similar predicted mechanism that might be preferable in terms of cost or FDA approval status.

As the application of explainable AI in the life sciences continues to grow, we anticipate that our framework will be broadly helpful to researchers. As observed in previous work, model prediction and model explanation are not always identical tasks [81, 82], and understanding how to create approaches that work for both of these goals is important given the popularity of Shapley value-based explanations for complex models. By demonstrating the high degree of variability in explanations within a class of models (Fig. 2-3), we hope to discourage users from naively selecting a single model to explain, and instead encourage users to explain ensembles of models. While our work focused on transcriptomic data, the high degree of feature correlation and dimensionality is also characteristic of many other forms of ‘omics data, indicating the broad impact of these results. We envision that future work on more efficient approaches to create ensemble models, which can be computationally costly, will be valuable. Likewise, further theoretical characterization of the feature attributions of complex models, such as deep neural networks and gradient boosting machines, will likely be important. While recent work has theoretically characterized the heterogeneity in feature importance across different well-performing models from the same model class, this work has thus far been limited to a small number of simple model classes (linear regression, logistic regression, and simple decision trees) [33].

In parallel to this work on improving the quality of attributions for black-box models, another thread of contemporary research focuses on incorporating prior biological knowledge into the modeling process. This includes methods like MERGE, which regularizes the coefficients of linear models using multiomic prior information [83], as well as Attribution Priors, which uses an efficient and axiomatic feature attribution method to align deep neural network attributions with biological priors during the training process [84, 85]. Other methods to incorporate biological prior information focus on structurally modifying neural network architectures, limiting interaction to genes that are known to share biological processes [86, 87]. Determining the best way to attribute feature importance in the context of the structurally-modified models will be important future work. Similarly, understanding how explainable AI can be optimally combined with the *unsupervised* deep learning models that have been successful in the context of single cell gene expression data will be another important line of future work [88, 89].

When applied to a large dataset of *ex vivo* drug synergy measurements in primary tumor cells from patients with AML, EXPRESS can both accurately predict drug synergy, as well as uncover a differentiation-related expression signature underlying the predictions for many combinations. While mutational status is increasingly considered in the clinical management of AML, our study demonstrates how useful tumor expression data can be for the prediction of drug combination synergy. Our experiments show that the extent of hematopoietic differentiation of AML cells is an important factor for the prediction of the synergy that can be achieved with specific therapy combinations, which has potential clinical application. One limitation of the current study is that our approach was applied to a dataset of drug synergy measurements in bulk tumor samples, rather than synergy assayed in specific purified tumor cell populations. As more and more studies come out specifically measuring the specific effects of anti-cancer drugs on the heterogeneous inidividual cells and subpopulations comprising AML [48, 66], applying EXPRESS to datasets with may yield interesting additional mechanistic insights.

## 4 Methods

### 4.1 Feature attribution methods

#### 4.1.1 Shapley values

The Shapley value is a concept from coalitional game theory designed to fairly distribute the total surplus or reward attained by a coalition of players to each player in that coalition [21]. For an arbitrary coalitional game, *v*(*S*) : *P*(*S*) *→*ℝ (where *S* is the set of players and 𝒫 indicates the powerset), the Shapley value for a player *i* is defined as the marginal contribution of that player averaged over the set of all *d*! possible orderings *R* of the *d* players in *S*:

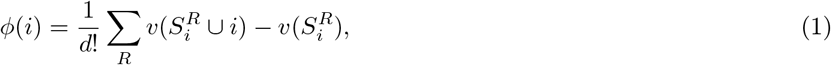

where 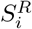 indicates the set of players in *S* preceding player *i* in order *R*.

To use this value to allocate credit to features in a machine learning model, the model must first be defined as a coalitional game. Deciding exactly how to define a model as a game is non-trivial, and a variety of different approaches have been suggested [17, 25, 90, 91]. The most popular, SHAP [17], defines the game as the conditional expectation of the output of a model *f* for a particular input sample *x* ∈ ℝ^*d*^ given that the features in *S* have been observed:

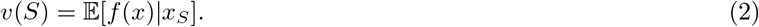

Because modeling an exponential number of arbitrary conditional distributions is often intractable, in practice the simplifying assumption that input features are independent is often made, allowing the expected value to be calculated over the marginal distributions of the features not in each given set, rather than the conditional distributions [17].

In our benchmark experiments, because comparable attributions are desirable for both the gradient boosted machine and neural network models, and because we want *global* attributions (features which are important for across all samples in the dataset), we use the SAGE software package to generate attributions. SAGE values define the coalitional game as the average reduction in test error *𝓁*(*·, ·*) when a set of features are included as compared to the base rate prediction *f*_ø_(*X*_ø_):

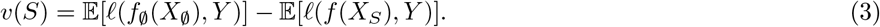

Since the SAGE package uses a sampling approach over possible coalitions of features to estimate Shapley values, it is important to ensure that the estimates are well-converged. To ensure convergence for the synthetic benchmark experiments, 102,400 permutations were used for all experiments (see Extended Data Fig. 6).

For experiments using the full BEAT AML dataset, we explained models using TreeSHAP [18]. TreeSHAP is a model-specific algorithm that leverages the structure of tree-based machine learning models (like XGBoost, the best performing model class for the problem) to quickly calculate SHAP values in polynomial time. TreeSHAP tries to approximate the conditional expectations using the conditional distribution defined by the tree structure. In instances where we needed global TreeSHAP attributions, we follow Lundberg et al. [18] and define the global attribution as the average magnitude of the local explanations *ϕ*_*i*_ over the whole dataset *D*:

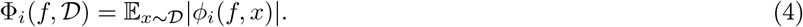

In instances where we wanted global attributions that were also directional, we considered the correlation between the SHAP attributions for a feature and that feature’s underlying value:

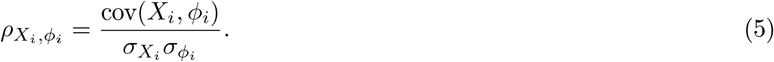

#### 4.1.2 Models used in benchmarks

In our benchmark tests, we evaluated two complex model classes explained using Shapley values. The first model class consisted of feed-forward neural networks. To train these networks, we used the PyTorch deep learning library [92]. To tune the models, we did a grid search across the following parameters: we used between 2 and 4 fully connected layers with either ‘ELU’ or ‘ReLU’ activations; we used a number either 64, 128, or 256 nodes in the first hidden layer and considered both a ‘decreasing’ and a ‘non-decreasing’ architecture (where ‘decreasing’ reduced the number of nodes in each successive layer by a factor of 2, and non-decreasing maintained a constant number of nodes across layers). We then trained the networks using the Adam optimizer with a learning rate of 0.001 for a maximum of 1,000 epochs. Early stopping was used to stop the training process if the mean squared error loss did not improve after 50 epochs. The second model class consisted of gradient boosted machines. To train these moodels, we used the XGBoost library [43]. To tune the models, we again did a grid search across several parameters: we considered a max tree depth of either 2, 10, 18, 26, 34, or 42; we also considered a range of ‘eta’ parameters including either 0.3, 0.2, 0.1, 0.05, 0.01, or 0.005. All models were boosted for 1000 rounds, and the saved model with best validation error was used for downstream prediction and explanation.

In addition to Shapley values applied to neural networks and gradient boosted machines, we also compared to a baseline of three more classical feature attribution methods used in biological feature discovery. The first involves ranking features *X* according to their Pearson correlation *ρ* with the outcome of interest *Y* :

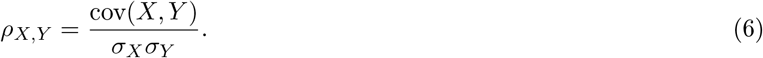

Ranking features in this way can be viewed as a special case of a family of feature selection algorithms known as Backward Elimination with the Hilbert-Schmidt independence criterion (BAHSIC) [35]. We also ranked features according to the magnitude of their coefficients in an elastic net regression, which is a linear regression where both the *𝓁*1 and *𝓁*2 norm of the coefficient vector are penalized in the loss function [36]. To train elastic net regression models, we used the ElasticNetCV function in the scikit-learn library with number of folds set to 5 [93]. Finally, we also tested a procedure known as recursive feature elimination using support vector machines (SVM RFE) [37]. As an estimator for this algorithm, we used the epsilon-Support Vector Regression function in scikit-learn with a linear kernel, then used the RFE function from the same library to select features with the parameters ’n_features_to_select’ and ’step’ set to 1.

### 4.2 Benchmark evaluation metric

To evaluate how well different approaches recover biologically-relevant signal, we designed a simple benchmark metric to evaluate the concordance between a list of features ranked by machine learning approaches and a ground truth list of features. It was necessary to design a new benchmark metric because existing metrics tend to evaluate how well feature attributions identify the features that are important for a particular machine learning model [18]. Our feature discovery benchmark measures how well each approach recovers biological signal by plotting the number of *true features* cumulatively found at each point in the list of features ranked by that approach, then summarizing this curve by measuring the area beneath it using the ‘auc’ function in scikit-learn [93] (see Extended Data Fig. 2). A larger area under the feature discovery curve (AUFDC) corresponds to better performance. A perfect score for a model with 10 true features out of 100 true features would be 950, while a random ordering would be expected to achieve an AUFDC of 500 on average. In order to make this score more intuitive, we subtract the random score of 500 and divide by the maximum possible area greater than random (450) so that the scores are scaled between 0 and 1, where 0 now means random performance and 1 means perfect performance.

### 4.3 Synthetic Datasets

In order to use our benchmark evaluation metric to determine how well different approaches could uncover underlying biological signal, it was essential to define datasets where the ground truth is known. Creating synthetic datasets also gave us the direct control needed to gain deeper understanding into the factors impacting the success of these algorithms, such as feature correlation, noise, and outcome type. We tested feature discovery performance on 240 total synthetic or semi-synthetic datasets. Each dataset comprised a feature matrix *X* ∈ ℝ^*n×d*^, where *n* represented the number of samples and *d* represented 100 input features, and an outcome vector *y ∈* ℝ^*n*^ which is some function of the original features (*y* = *f* (*X*)).

We considered three groups of distributions for the feature matrices. The first group was 1000 samples of 100 independent Gaussian features randomly generated to be 0 mean, unit variance. The second group was 1000 samples of 100 Gaussian features with ten groups of five tightly correlated features (Pearson’s *ρ* = 0.99). The final group involved 223 real patient gene expression samples from the Beat AML Dataset [7].

We considered four different functions *f* by which the features *X* were related to the outcome *y*. The first function was a linear function with 10 non-zero coefficients, *f* (*X*) = *Xβ*. The second function was the sum of 10 univariate ReLU functions 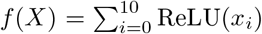. The third function was a sum of 10 pairwise multiplicative interactions 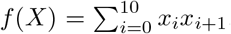. The final function was a sum of 10 pairwise AND functions 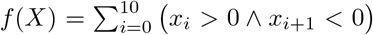. For each of the twelve possible pairwise combinations of feature matrices and outcome functions, we created 20 specific datasets meeting the specifications, where the only difference was that the features were randomly regenerated (or randomly re-sampled from the full transcriptome in the case of the AML features), and the features selected as true features were re-selected.

### 4.4 Comparing ensembles and individual models

To train ensemble models for comparison in our benchmark experiments, we used the method of bootstrap aggregation, or *bagging* [94]. This method involves first bootstrapping the data, or resampling the dataset with replacement until the bootstrapped dataset has as many samples as the original, then training a model on the bootstrap resampled dataset. We repeat the process of bootstrapping and training models 20 times. Since our benchmark is a regression problem, the 20 model outputs are then aggregated by a simple mean. This method is known to improve predictors by increasing their stability.

To understand the difference in quality of the individual model attributions and the ensemble model attributions, we considered two separate objective metrics. The first was to assess the *stability* of the attributions. We measured the pairwise cosine similarity of 20 ensemble models’ attributions trained on bootstrap resampled versions of each dataset, then measured the pairwise cosine similarity of 20 individual models’ attributions trained on bootstrap resampled versions of the same datasets:

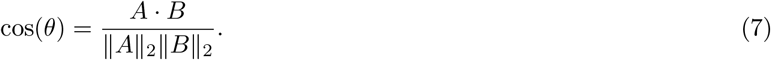

The next metric aimed to understand how much importance was put on truly important features compared to how much was potentially placed on spurious correlates. We therefore measured the Gini index of each global attribution vector to understand how *sparse* of an attribution was learned by each model:

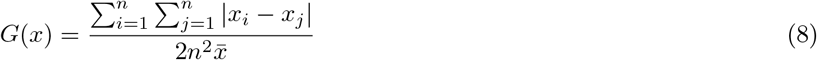

### 4.5 Beat AML Datasets

The Beat AML program comprises a large cohort of AML patient tumor samples for which *ex vivo* anti-cancer drug sensitivity has been measured. Since our project aimed to uncover the transcriptomic factors underlying anti-cancer drug synergy, we only included patients from the cohort whose tumors had been characterized with RNA sequencing, for which measurements pairs of anti-cancer drugs had been tested. Our final dataset contained the RNA-sequencing expression data from 285 patients with myeloid malignancy, and drug synergy measured on a subset of patients for 131 combinations of 46 distinct drugs.

The input features used in modeling each of 12,362 samples (where a sample is one patient and one combination of two anti-cancer drugs) were represented as a vector *x ∈* ℝ^15535^. This vector is constructed by concatenating three other vectors. First, we describe each patient’s tumor sample using a vector of gene expression values (RNA-seq data – see RNA-seq pre-processing section for more information), *g ∈* ℝ^15377^. We described each drug combination using a feature vector, *v∈*ℝ^46^, of drug identity labels where each element *v*_*i*_ was equal to 1 if the *i*th drug was present in the combination and 0 otherwise. We also incorporated drug target information for each drug combination, using information compiled from DrugBank plus a supplementary literature search for reliable drug targets, for a total set of 146 targets.1 We then described the drug targets of each combo with a vector *u ∈* ℝ^146^, where each element *u*_*j*_ was equal to 2 if the *j*th target was targeted by both drugs, equal to 1 if the *j*th target was targeted by only one of the drugs, and equal to 0 if the *j*th target was not targeted by either drug.

### 4.6 RNA-seq preprocessing

To ensure a quality signal for prediction while removing noise and batch effects, it is necessary to carefully preprocess the RNA-seq gene expression data. In this study, the RNA-seq were preprocessed as follows. First, raw transcript counts were converted to fragments per kilobase of exon model per million mapped reads (FPKM). FPKM is a more reflective of the molar amount of a transcript in the original sample than raw counts, as it normalizes the counts for different RNA lengths and for the total number of reads.6 FPKM is calculated as follows:

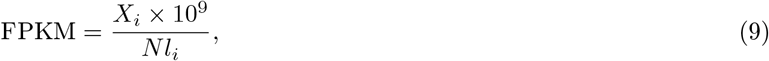

where *X*_*i*_ represents the raw counts for a transcript, *l*_*i*_ is the effective length of the transcript, and *N* is the total number of counts.

After converting counts to FPKM, we removed any non-protein-coding transcripts from the dataset. We also removed transcripts that were not meaningfully observed in our dataset by dropping any transcript where *>* 70% measurements across all samples were equal to 0. We then log-transformed the data and standardized each transcript across all samples, such that the mean for that transcript was equal to zero and the variance of the transcript was equal to one. Finally, we corrected for batch effects in the measurements using the ComBat tool available in the sva R package [95].

### 4.7 Drug synergy metric

The outcome in our model was drug synergy: whether a number of drugs exhibit more anti-cancer activity in combination than would be expected simply by adding their individual activities together. We therefore calculated synergy using the Combination Index (CI) of the two drugs:

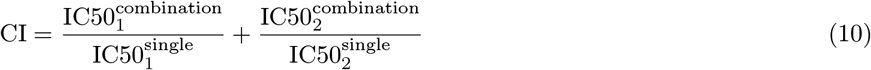

where 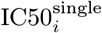 is the dose of drug *i* required to reduce cell viability to 50% when used alone and 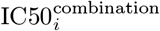 is the dose of drug i required to reduce cell viability to 50% when used in combination with the other drug we are measuring [96]. When a drug combination is synergistic, the CI will be less than 1 (it will be equal to 1 when the combination is additive and greater than 1 when the combination is antagonistic). In our model, we log-transformed the CI measure to help manage the skewness of the original distribution, and then scaled the measure to make the distribution 0 mean and unit variance. We also multiplied by -1 for ease of interpretation: more synergistic combinations thus have a larger score.

While prior studies have made use of response-surface analysis, which involves measuring the volume between an idealized additive response surface and a measured actual response surface (9,10), these measures could not be applied to the “diagonal” measurements present in the Beat AML Dataset. A major drawback to response-surface analyses is they requires a “checkerboard” of measurements at different drug concentrations, where the ratios and doses of each drug in a combination is varied. This consumes many more cancer cells, which is problematic when using primary cells from patients, as the amount of sample that can be collected is more limited than when using cell lines.

### 4.8 Cross-validation and sample stratification

In addition to the model parameters which are learned from data, machine learning models also rely on hyperparameters, which must be tuned to a specific task in question in order to attain optimal predictive performance. In order to estimate the true generalization error of a model (i.e., how well that model is likely to perform on unseen data), it is essential that model parameters and hyperparameters must be learned and chosen based on training data, while predictive performance is evaluated on a held-out test set that is never used for hyperparameter selection or model training. Hyperparameters are typically picked through a cross-validation (CV) procedure which determines the optimal hyperparameters for the model by validating them through a number of internal training and validation fold pairs randomly chosen from the set of training samples used for learning the model parameters.

To effectively train our models and evaluate predictive performance, we therefore utilized a nested 5-fold CV procedure, whereby the data were split into 5 separate test folds. For each of these test folds, we trained our synergy prediction model using the four remaining folds and evaluated it on the held-out test fold. To properly tune the hyperparameters of the models trained for each test fold, three of the four training folds were used as an internal training set, while the remaining fold was used as a validation set. The hyperparameters were selected by an inner loop, where for each hyperparameter set of interest, the model was trained on the internal training set and tested on the validation set. The hyperparameters giving the best performance on the validation set were then used to train a model on the entire training data, which was then finally evaluated on the held-out test fold. The grid of hyperparameters tested for each model type are as follows. For the sklearn elastic net implementation, the “alpha” parameter was tuned over values ranging from 0.1 to 100, while the “l1_ratio” parameter was tuned from 0.25 to 0.75. For the sklearn Random Forest implementation, the “n_estimators” parameter was tuned from 128 trees to 2048 trees, while the “max_features” parameter was set to be either “log2,” “sqrt,” or “256.” For XGBoost, “max_depth” was tuned between 4 and 8, “subsample” was tuned to values between 0.1 and 0.8, and learning rate was tuned between 0.05 and 0.1. For deep neural networks, hyperparameters were tuned following the grid given in [14], where an additional data pre-processing step that would optionally transform the RNA-seq features with a hyperbolic tangent function in addition to standardization was also included as a hyperparameter. For both deep neural networks and gradient boosted machines, early stopping based on validation set error was used to choose the number of epochs/estimators.

In order to evaluate the model’s performance for a variety of hypothetical uses, we stratified our data into training and testing sets in four different ways (Fig. 4). Each sample in our dataset consists of a synergy measurement for a 2-drug combination tested in a patient’s tumor cells. In the first stratification setting, we ensured that any sample (2-drug combination and patient) present in the test data would never be present in the training data. The second setting maintains the first setting’s requirement that each sample in the test data be novel, but additionally ensures that any combination of drugs in the test data would never be present in the training data. The third setting maintains the first setting’s requirement that each sample in the test data be novel, but additionally ensures that any patient in the test data would never be present in the training data. Finally, the fourth setting maintains the first and second settings’ requirements, while additionally ensuring that for any combination of drugs in the test data, at least one of the drugs in that combination would never have been present in the training data. Each of these settings should be increasingly difficult to predict, as each setting requires progressively more generalizable trends in the data to have been learned.

### 4.9 XGBoost model ensembles

After selecting XGBoost as the best-performing model class for the prediction of anti-AML drug synergy, we then wanted to account for the full diversity of possible good XGBoost models fit to the highly correlated AML gene expression data. We therefore trained 100 models and explained the ensemble model. Each individual model had both row and column subsampling turned on for each additional tree fit, and the difference between the models in the ensemble was the random seed given to generate the subsampling.

In practice, instead of explaining the entire ensemble (the average output of each of the 100 models), we instead explain each individual model and average the explanations. This is possible due to the linearity property of Shapley values [97]. This property states that for the convex combination of any two coalitional games *v* and *w*, the attribution for player *i* will be the convex combination of the attributions that player would attain in each individual game *v* and *w*:

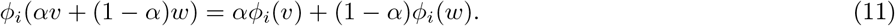

### 4.10 Overall pathway analysis

The highest-ranked genes in the lists ordered by global Shapley values were tested for pathway enrichments using the StringDB package in R [98]. The package was initialized using the arguments: ‘version’ = ‘10’, ‘species’ = ‘9606’, and ‘score_threshold’ set to the default of 400. We used the set of pathways from KEGG (Kyoto Encyclopedia of Genes and Genomes) for enrichment tests. The actual enrichments were calculated by a hypergeometric test implemented in the ‘get_enrichment’ method. In order to ensure that pathway enrichments were robust to the threshold used for selecting the highest-ranked genes, we averaged the enrichment test result over a variety of different thresholds, ranging from 200 to 800 top genes. FDR correction was applied using the Benjamini-Hochberg procedure [99].

### 4.11 Generation of Differential Expression Stemness Profiles and Measurement of Synergy Correlation

To generate expression signatures related to more or less-differentiated states of cells in the hematopoietic cell lineage, we downloaded RNA-seq data from isolated cells from particular levels of developmental transitions [69]. We then used the R package DESeq2, which tests for differential expression in RNA-seq data based on a negative binomial model, to generate lists of genes upregulated in particular populations of cells as compared to other populations [100]. The populations we compared were as follows: monocytes vs. hematopoietic stem cells (HSCs), lymphocytes vs. HSCs, all fully differentiated cells (which included erythroblasts, T cells, B cells, NK cells, and monocytes vs. HSCs, monocytes vs. leukemic stem cells (LSCs), lymphocytes vs. LSCs, and all fully differentiated cells vs. LSCs. The immunophenotypes used to sort HSCs and LSCs were Lin– CD34+ CD38– CD90+ CD10– and Lin– CD34+ CD38– TIM3+ CD99+, respectively. Gene expression profiles for these populations were the same used in [69]. The multiple testing-adjusted p-value used as a significance threshold for differentially-regulated genes was 0.05.

When we tested for association between our differential expression profiles and synergy for particular combinations of drugs, we first considered only samples containing the drug combination in question. We then averaged gene expression over all genes in the differential expression profile. Finally, we measure the Pearson correlation between the average expression profile and the drug combination synergy for those samples. Since we have many combinations of drugs and many differential expression profiles to test, we correct for multiple testing using the Benjamini-Hochberg FDR correction procedure [99]. We then display only correlations that are significant after correction. Additionally, we only want to consider correlations that are robust to the differences in the particular stemness-differentiation profiles, so we only plot correlations for drugs that are significant across at least two profiles.

### 4.12 Drug-specific pathway analysis

To analyze the biological processes relevant for combinations containing specific drugs in the dataset, we tested the top-ranked genes in the lists ordered by the average magnitude Shapley interaction indices [18, 71]. Following the same procedure described above, we calculated pathway enrichments using the StringDB package in R [98]. We used the set of pathways from Gene Ontology (GO) Biological Process terms for enrichment tests. The enrichments were calculated by a hypergeometric test implemented in the ‘get_enrichment’ method. In order to ensure that pathway enrichments were robust to the threshold used for selecting the highest-ranked genes, we averaged the enrichment test results over a variety of different thresholds, ranging from 200 to 800 top genes. FDR correction was applied using the Benjamini-Hochberg procedure [99].

For Venetoclax, since a large number of biological process terms were signficicantly enriched, and since there is significant overlap and similarity between these gene sets, we clustered the significantly enriched pathways into modules. We defined an adjacency matrix where each gene set represented a node in a network, and the Jaccard Index (a measure of overlap) between pathways was used to define edges. We binarized the matrix for pathways with Jaccard Index greater than 0.4. We then manually annotated all connected components in the resultant graph (see Supplemental Table 6). To plot the network, we used the spring layout functionality in the networkx library in Python [101].

## 5 Data availability

The results published here are in part based upon data generated by the Cancer Target Discovery and Development (CTD2) Network (https://ocg.cancer.gov/programs/ctd2/data-portal) established by the National Cancer Institute’s Office of Cancer Genomics. Sequencing data are available in the GDC data portal under dbGaP Study Accession phs001657. The Beat AML patient sample data used in this study was done under an early access agreement, prior to final accrual, harmonization, and public release of the full dataset. As such, the subset of samples included in this study may differ in sample representation, quality control thresholds, and data normalizations from those found in GDC and in the final study describing the full dataset.

The code necessary to reproduce synthetic datasets can be found at https://github.com/suinleelab/express.

## 6 Code availability

Code necessary to reproduce our experimental findings can be found at https://github.com/suinleelab/express.

## 8 Materials & Correspondence

Correspondence to Su-In Lee and Kamila Naxerova.

## Extended Data

**Extended Data Fig. 1.**
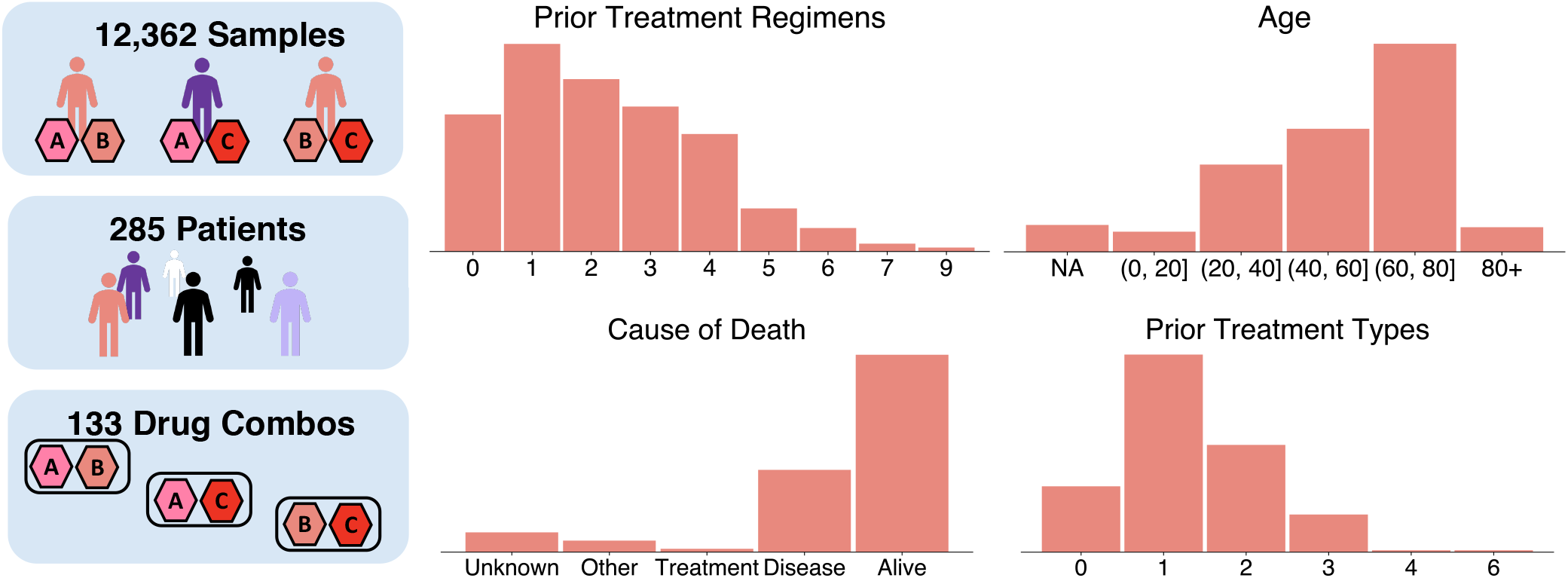
Descriptive statistics of Beat AML cohort. Histograms showing the relative density of prior treatment regimens, age, cause of death, and prior treatment types in the cohort of 285 patients in our dataset, which consisted of 12,362 samples with paired gene expression and drug synergy measurements for 133 pairs of 46 anti-cancer drugs.

**Extended Data Fig. 2.**
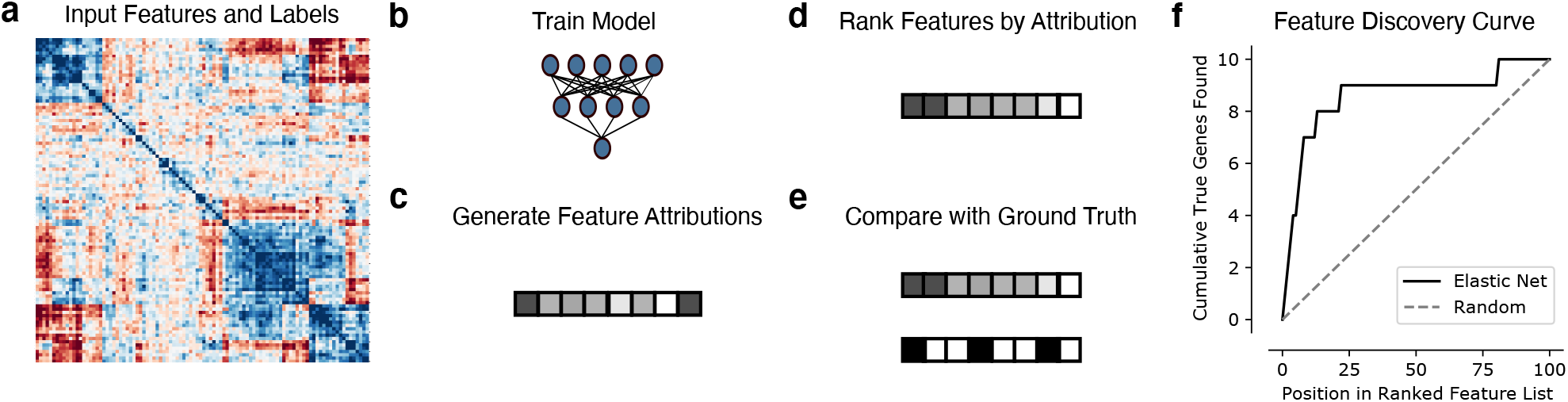
Feature discovery benchmark. For each synthetic or semi-synthetic dataset (**a**), we trained a variety of models (**b**) including neural networks, gradient boosted machines, support vector machines, and elastic net regression, as well as univariate statistics (Pearson correlation). For the machine learning models, we then used SAGE to generate global Shapley value feature attributions (**c**), ranked the features according to the magnitude of their attributions (**d**), and compared the ranked list generated by each method to the binary ground truth importance vector (**e**). To measure the feature discovery quality of each method, we plotted how many “true” features are found cumulatively at each point in the ranked feature list (**f**), then summarized the curve generated by this procedure by measuring the area under the feature discovery curve (AUFDC). This score is then rescaled so that a score of 0 represents random performance while a score of 1 represents perfect performance.

**Extended Data Fig. 3.**
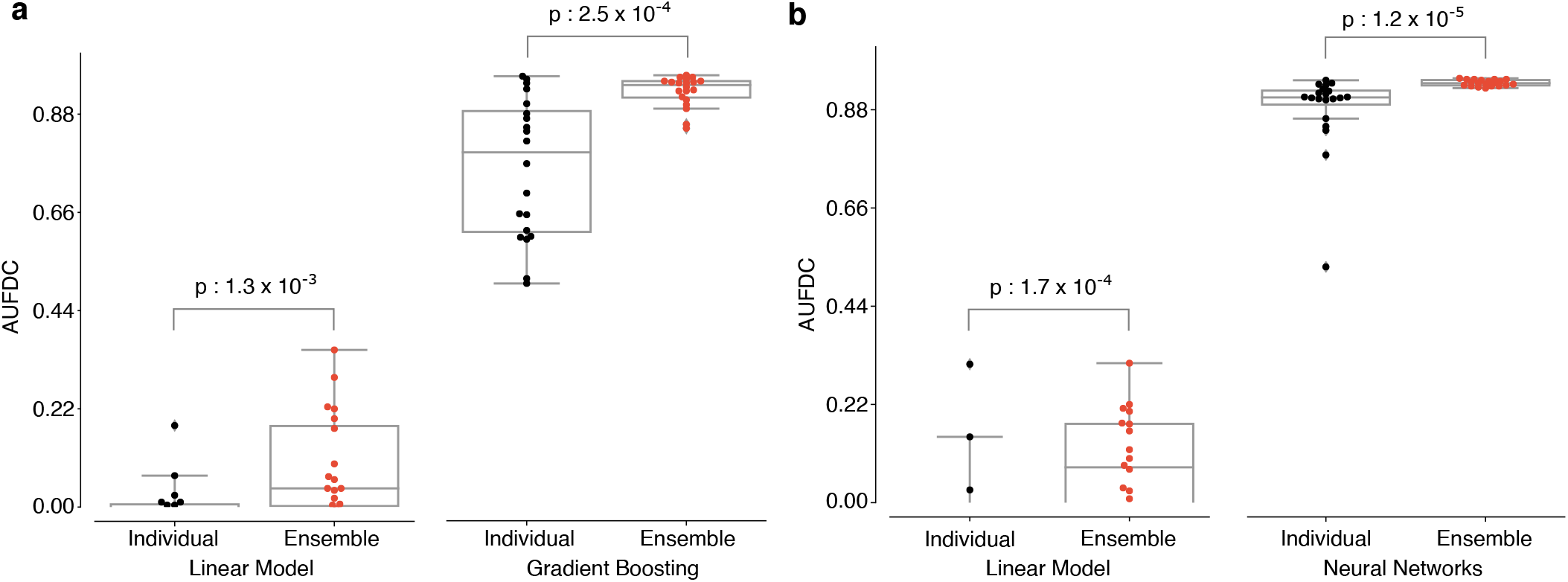
Explaining ensembles overcomes variability on correlated groups dataset. **a-b**, Comparison of feature discovery performance between individual models and ensemble models on the correlated groups dataset with a step function outcome (**a**) and a multiplicative outcome (**b**).

**Extended Data Fig. 4.**
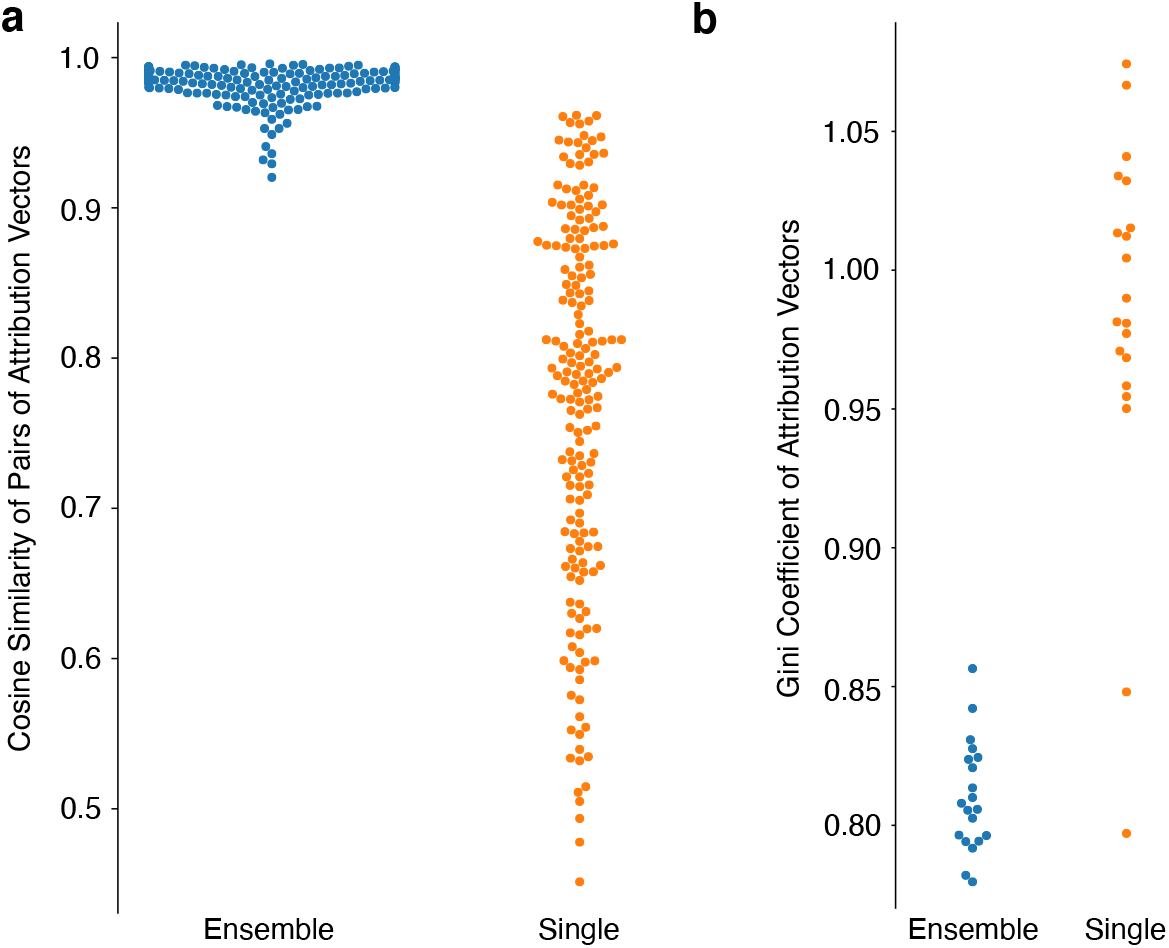
Synthetic data attribution characteristics. To understand why ensemble models were able to attain better feature discovery performance than single models, we compared the characteristics of the attribution vectors of XGBoost models trained on bootstrap resampled versions of a Correlated Groups dataset with a step-function outcome. **a**, Pairs of attribution vectors from ensembled models are more similar across bootstrap resamples of the dataset than attribution vectors from single models, as measured by cosine similarity. **b**, Attribution vectors from ensembled models place a larger proportion of their importance on a smaller set of features than attribution vectors from single models, as measured by the gini coefficient of the attribution vectors, a measure of vector sparseness.

**Extended Data Fig. 5.**
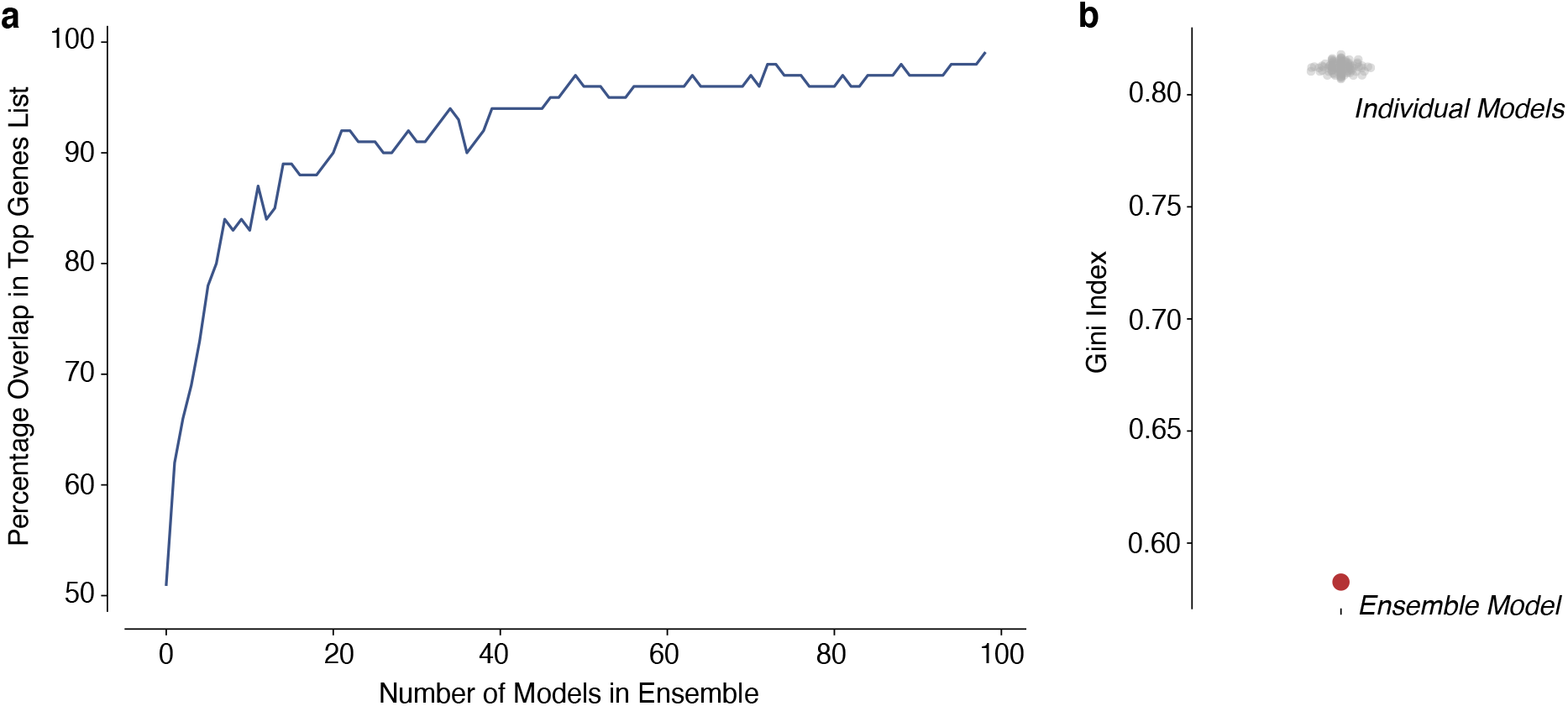
Beat AML data attribution characteristics. **a,** To ensure that a sufficient number of models were included in our final ensemble, we measured the percentage overlap in the final list of top 100 genes, and the cumulative top 100 genes list as additional XGBoost models were ensembled. **b** Attribution vector sparseness for individual models trained on Beat AML dataset (grey) and attribution vector sparseness for our final ensemble model (red). A lower Gini Index indicates a more sparse attribution vector.

**Extended Data Fig. 6.**
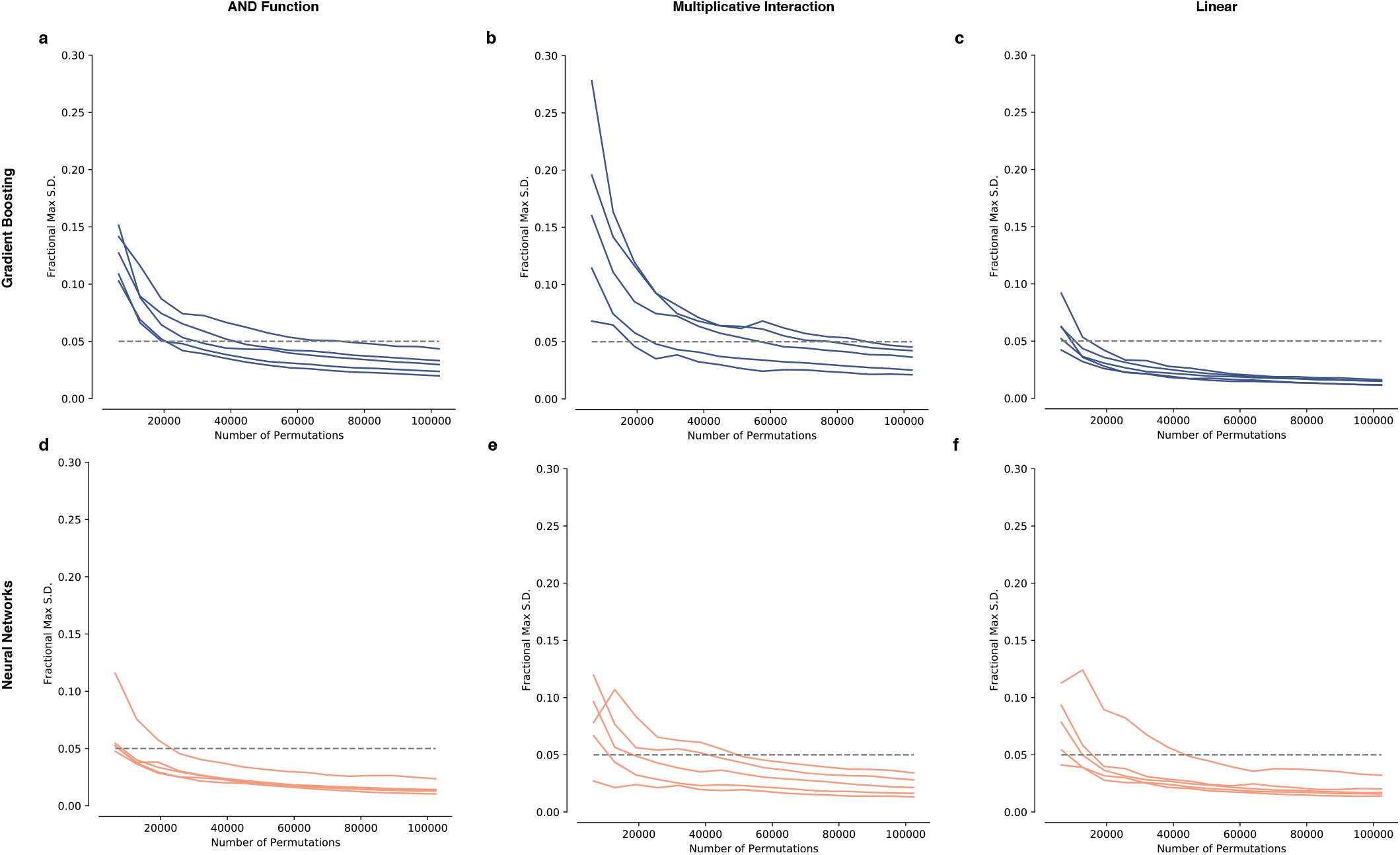
Sampling convergence of SAGE values. To ensure convergence of the Shapley value estimates used in our benchmark, we measured the maximum standard deviation of the elements in our attribution vector (generated using SAGE) as a fraction of the total attribution (Fractional Max S.D.), plotted against the number of permutations sampled. We found that for all three outcome types (a step function generated using a boolean AND function, a sum of pairwise multiplicative interactions, and a linear function), for both gradient boosting models (**a-c**) and neural networks (**d-f**), 102400 permutations were sufficient to drive the Fractional Max S.D. below 5%.

